# GLP-1 Receptor Agonist Improves Metabolic Disease in a Pre-clinical Model of Lipodystrophy

**DOI:** 10.1101/2023.09.01.555852

**Authors:** Ahlima Roumane, George D. Mcilroy, Nadine Sommer, Weiping Han, Lora K. Heisler, Justin J. Rochford

## Abstract

Individuals with lipodystrophies typically suffer from significant metabolic disease linked to adipose tissue dysfunction including severe insulin resistance and lipoatrophic diabetes, hepatic steatosis and hyperphagia. Current treatment options are limited and beter therapies for affected individuals are urgently needed. No systematic, detailed analyses exist of the effects of glucagon like peptide-1 receptor (GLP-1R) agonists in the treatment of lipoatrophic diabetes. Here we examined the effects of the GLP-1R agonist liraglutide in seipin knockout mice, a pre-clinical model of generalised lipodystrophy. Acute liraglutide treatment of seipin knockout mice significantly improved insulin, glucose and pyruvate tolerance. Once-daily injection of seipin knockout mice with liraglutide for 14 days led to a modest reduction in food intake but significant improvements in hepatomegaly associated with steatosis and significantly reduced markers of liver fibrosis. Detailed examination of the pancreas revealed that liraglutide enhanced insulin secretion in response to glucose challenge with concomitantly improved glucose control. Thus, GLP-1R agonist liraglutide significantly improved multiple aspects of lipoatrophic diabetes and hepatic steatosis in mice with congenital generalised lipodystrophy. This provides important insights regarding the benefits of GLP-1R agonists for treating lipodystrophy, informing more widespread use to improve the health of individuals with this condition.

## Introduction

Syndromes of lipodystrophy are rare disorders where reduced adipose mass or adipose tissue dysfunction leads to metabolic disease. Affected individuals are typically characterised by lipoatrophic diabetes, insulin resistance and hepatic steatosis with a complex array of additional features making this a multisystem disease [1-5]. Although rare, lipodystrophies are likely to be underdiagnosed [6]. In congenital generalised lipodystrophies (CGL) adipose tissue may be almost absent. In more frequently occurring familial partial lipodystrophy (FPLD) affected individuals may have modest changes in absolute adipose mass but redistribution or dysfunction of adipose tissues [1; 2]. Metreleptin is the only specific treatment to alleviate the metabolic complications of lipodystrophy. It is highly effective in CGL but is only effective in a subset of individuals with FPLD, typically those with low circulating leptin levels [7-9]. Moreover, metreleptin requires daily injection, which is painful in lipodystrophic individuals due to the paucity of subcutaneous adipose tissue, is costly and not available to patients in all countries. Hence, there is an urgent need for new treatments.

Agonists of the glucagon-like peptide-1 receptor (GLP-1R) deliver significant beneficial effects in treating type 2 diabetes and obesity [10; 11]. Key effects of GLP-1R agonists are to produce improvements in insulin sensitivity, reduce food intake, reduce adiposity, and delay gastric emptying [10; 11]. Lipoatrophic diabetes observed in patients with lipodystrophies differs significantly from common obesity-associated type 2 diabetes in several regards, typically with significantly reduced adipose mass, dysregulation of adipokines, greater severity of dyslipidaemia and insulin resistance and hyperphagia [1; 2; 4]. There are currently no systematic analyses of the effects of GLP-1R agonists in lipodystrophy, only a very small number of case reports [12-14]. Hence, there remains no detailed information regarding the effects of GLP-1R agonists specifically in lipoatrophic diabetes. This is critical to obtain before widespread adoption of these drugs in individuals with lipodystrophy. To address this, we used a well validated *Bscl2*-null (seipin knockout) mouse model of CGL to define the effects of GLP-1R agonists and inform the adoption of this class of drugs for individuals with lipodystrophy.

## Materials and Methods

### Animals

Seipin knockout (SKO) *Bscl2*-null mice were as previously described in [15]. All procedures were carried out in accordance with the U.K. Animals (Scientific Procedures) Act 1986 and following local ethical approval at University of Aberdeen. Unless otherwise stated, mice had *ad libitum* access to water and standard rodent chow diet (CRM (P) 801722, Special Diets Services).

### Tolerance tests and drug treatment

Mice (22- to 25-week-old) were fasted for five hours and intraperitoneal (i.p.) injections of 0.2 mg/kg liraglutide (Tocris) or phosphate-buffered saline (PBS) were 45 minutes before the tolerance tests (-45 min). Glucose levels were determined by glucometer readings (AlphaTrak® II, Zoetisus) from tail punctures. Mice received i.p. injection of 2 mg/g D-Glucose (Sigma-Aldrich) for glucose tolerance tests (GTT), 0.75 IU/kg of human insulin (Actrapid 100 IU/mL, Novo Nordisk) for insulin tolerance tests (ITT) or 2 g/kg pyruvate (Sigma-Aldrich) for pyruvate tolerance tests (PTT) as in [16]. Blood glucose levels were monitored at 15, 30, 60, 90 and 120 minutes. Alternatively, mice received a daily i.p. injections of either 0.2 mg/kg liraglutide (Tocris) or vehicle (PBS) for 14 days.

### Metabolic assessment

Mice were singly housed in cages automatically collecting food and water intake data (PhenoMaster, TSE Systems). Mice acclimatized to the cages for 3 days prior to measurements. Food intake and water consumption were measured over 8 consecutive days as in [16].

### Serum analysis

Blood was collected by cardiac puncture from five-hour fasted mice and serum isolated in SST™ Amber tubes (BD Microtainer®). Serum triglyceride contents were determined using the Triglyceride Liquid Assay (Sentinel Diagnostics) following manufacturer’s instructions. Serum insulin levels were measured by ELISA (Mercodia).

### Tissue and histological analyses

Triglyceride levels were determined in supernatants prepared from frozen liver samples using the Triglyceride Liquid Assay (Sentinel Diagnostics) as in [17]. Liver tissue and pancreata were fixed in 10% neutral buffered formalin, paraffin-embedded and 5 µm-thick sections were stained with hematoxylin and eosin (H&E) or picrosirius red solution (Abcam #ab150681) as in [18]. For insulin staining, pancreatic sections were incubated overnight at 4°C with anti-insulin antibody (1:500, Clone L6B10, Cell Signaling) and detection with an HRP-conjugated secondary antibody using 3,3’-diaminobenzidine (DAB, Vector Laboratories, #SK-4100) substrate. Between 6 and 10 pancreata per gender and genotype and for each treatment were analysed for β-cell area quantification. For each pancreas, 3 sections 150 μm apart were analysed and total area occupied by insulin-positive cells determined using QuPath sotiware [19]. RNA extraction and real-time quantitative PCR was carried out as in [17]. Gene expression was normalised to the geometric mean of three stable reference genes (*Nono, Ywhaz* and *Hprt*). Sequences and details of primers and assay probes are provided in the Supplementary Table S2.

### Statistical analyses

All the data are presented as mean ± SEM and analysed by unpaired two-tailed Student’s t-test or two-way ANOVA with Bonferroni or Tukey’s post-hoc test as appropriate using GraphPad Prism. P < 0.05 was considered as statistically significant.

## 3. RESULTS

### 3.1 Liraglutide improves insulin and glucose tolerance in lipodystrophic mice

Seipin knockout (SKO) mice were injected with PBS or Liraglutide (0.2 mg/kg) 45 min before performing insulin, glucose, and pyruvate tolerance tests (ITT, GTT and PTT respectively). SKO mice failed to respond to insulin during an ITT, consistent with severe insulin resistance (Fig. 1 A, C). Liraglutide treatment significantly improved insulin sensitivity in SKO mice compared to PBS-treated SKO controls (Fig. 1A-D). SKO mice also showed impaired glucose tolerance compared to WT mice (Fig. 1E, G). Glucose clearance rates were significantly improved in liraglutide-treated SKO mice and in females were similar to those observed in liraglutide-treated WT mice (Fig. 1E-H). In a PTT, pyruvate injection led to significantly greater increases in blood glucose levels in SKO *versus* WT mice injected with PBS, suggesting poorer control of gluconeogenesis. Liraglutide treatment of female SKO mice reduced blood glucose levels below those observed in WT PBS-treated mice (Fig. 1I, J). This effect was more modest in male SKO mice and failed to reach statistical significance (Fig. 1K, L). Collectively, these data show that acute liraglutide treatment significantly improves glucose and insulin tolerance, and control of hepatic gluconeogenesis in SKO mice, most strikingly in female SKO mice.

**Figure 1:**
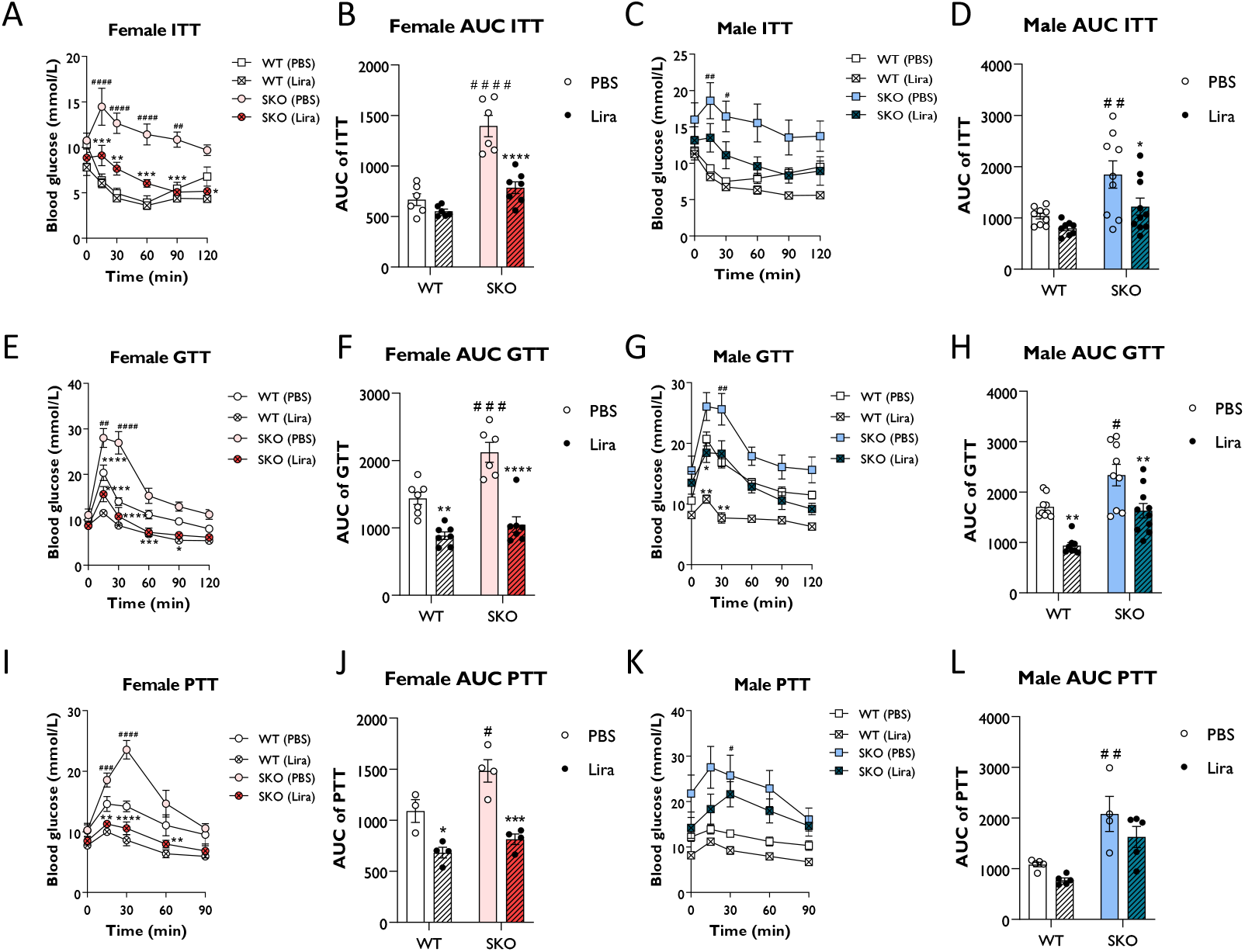
Liraglutide improves glucose tolerance and insulin sensitivity in mice with lipodystrophy. Male or female 22-week-old wild-type (WT) and seipin knockout (SKO) mice were intraperitoneally injected with phosphate-buffered saline (PBS) or 0.2 mg/kg liraglutide (Lira) prior to insulin tolerance tests (ITT) (**A-D**), glucose tolerance tests (GTT) (**E-H**) and pyruvate tolerance tests (PTT) (**I-L**). Areas under the curve (AUC) are shown for ITTs (**B, D**), GTTs (**F, H**) and PTTs (**J, L**) in female and male mice as indicated (female n= 3-7, male n= 4-10). Data are presented as mean ± SEM; *p < 0.05, **p < 0.01, ***p < 0.001, ****p < 0.0001 (PBS versus liraglutide within same genotype); ^#^p < 0.05, ^##^p < 0.01, ^###^p < 0.001, ^####^p < 0.0001 (WT (PBS) versus SKO (PBS)).

### 3.2 Effect of liraglutide on food and water intake, body composition and glycaemia

To examine longer-term effects of liraglutide WT and SKO mice were given daily injections of PBS or 0.2 mg/kg liraglutide. Food and water intake was measured between days 4 and 12. As observed in CGL2 patients, SKO mice were hyperphagic (Fig. 2A), atributable to the low circulating leptin levels [20; 21]. Water intake was also significantly elevated in SKO mice (Fig. 2B). Liraglutide significantly decreased cumulative food intake in male SKO mice, with a small, non-significant reduction observed in females (Fig. 2C). Female and male SKO mice injected with liraglutide showed a progressive normalisation of water consumption with a 45% and 62% reduction, respectively, by day 12 (Fig. 2D). This likely reflects an anti-diabetic treatment effect of liraglutide reducing polydipsia in SKO mice. As shown previously [15; 20], lipodystrophic SKO mice display significantly reduced fat mass (Fig. 2E) and increased lean mass (Fig. 2F) relative to WT mice. Liraglutide significantly lowered fat mass in male WT mice, but not in female WT mice (Fig. 2E) whilst lean mas was significantly increased in both male and female WT mice (Fig. 2F). In contrast, body composition of SKO mice was not affected by liraglutide or vehicle treatment (Fig. 2E, F). Liraglutide-treatment for 12 days did not alter non-fasted blood glucose levels in WT mice, which remained around 10 mmol/L. Prior to treatment, SKO mice were hyperglycaemic (Fig. 2G). Liraglutide treatment significantly reduced hyperglycaemia in both male and female SKO mice with no effect observed in PBS-treated mice (Fig. 2G).

**Figure 2:**
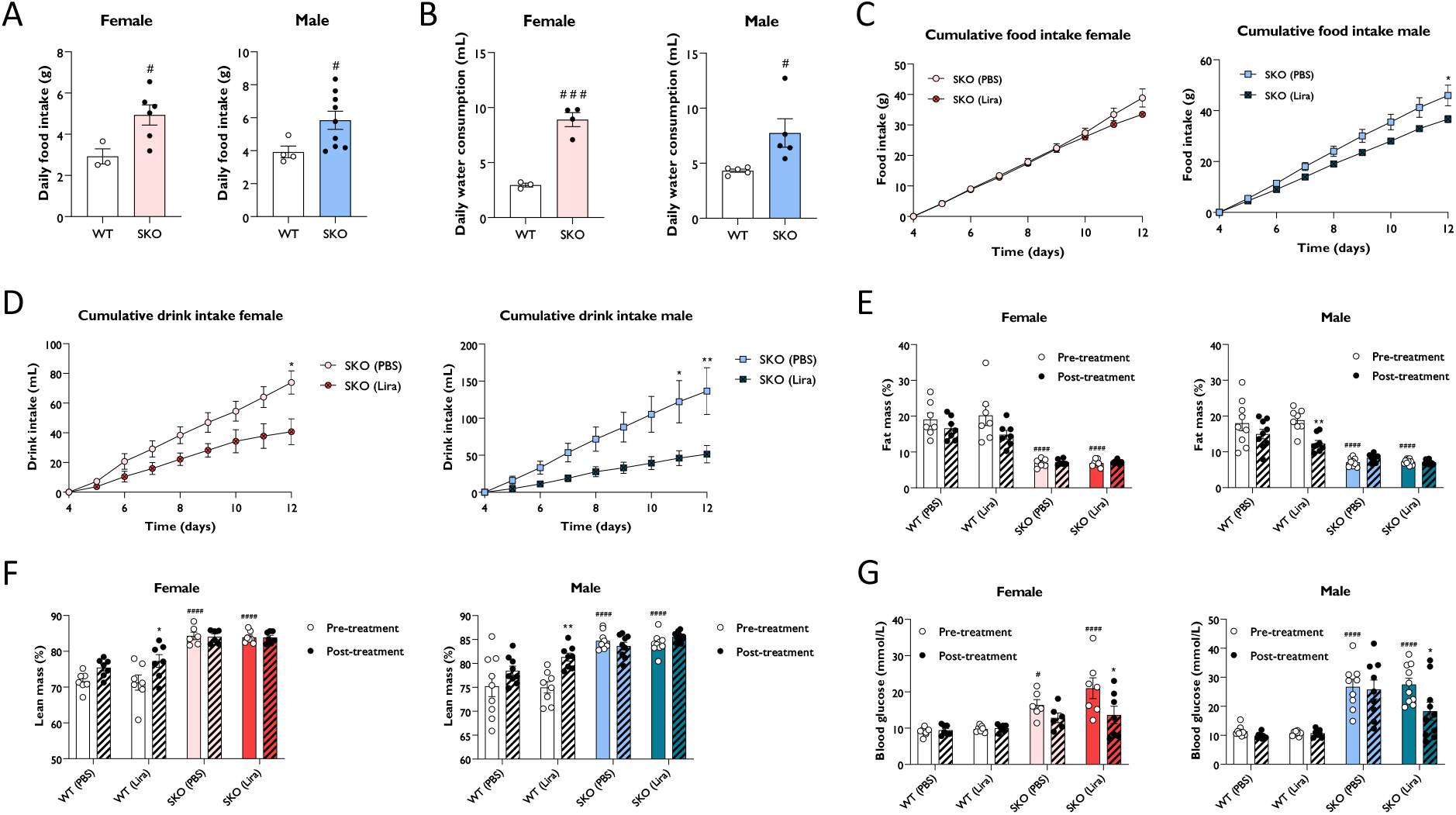
Effect of liraglutide on food and water intake, body composition and glycaemia. Average daily food intake (**A**) and water consumption (**B**) for 22-week-old singly-housed wild-type (WT) and seipin knockout (SKO) mice measured over three to seven days. Female and male WT and SKO mice were intraperitoneally injected once daily with phosphate-buffered saline (PBS) or 0.2 mg/kg liraglutide (Lira) as indicated. Food intake (**C**) and water consumption (**D**) were measured from day 4 to day 12 of treatment. Fat mass (**E**), lean mass (**F**) and non-fasted blood glucose levels (**G**) were determined on day 12 (female n= 3-7, male n= 4-10). Data are presented as mean ± SEM; *p < 0.05, **p < 0.01 (PBS versus liraglutide within same genotype); ^#^p < 0.05, ^###^p < 0.001, ^####^p < 0.0001 (WT (PBS) versus SKO (PBS)).

### 3.3 Liraglutide improves hepatic steatosis and fibrosis development in lipodystrophic mice

Consistent with the hepatic steatosis seen in individuals with CGL [20; 22; 23], SKO mice displayed significantly enlarged livers when compared to WT controls with substantial accumulation of triglycerides (Fig. 3A, B). Daily liraglutide treatment for 14 days significantly decreased liver weights by 24% in male and 27% in female SKO mice (Fig. 3A). Hepatic triglyceride content was also reduced in female SKO mice not significantly different between PBS and liraglutide-treated male SKO mice (Fig. 3B). H&E staining of liver sections also revealed significant lipid accumulation in SKO mice that was considerably reduced by liraglutide treatment in female but not male SKO mice (Fig. 3C, D). Picrosirius red staining showed substantial fibrosis in both male and female SKO mice but not WT mice which was reduced by liraglutide treatment (Fig. 3C, D). In agreement with histological staining, mRNA levels of several pro-fibrogenic markers, including *Mmp13, Timp1, Tgfβ* and *Col3a1* were significantly upregulated in both male and female PBS-treated SKO mice *versus* PBS-treated WT mice (Fig. 3E, F). Liraglutide treatment significantly reduced *Mmp13, Timp1, PAI-1, Tgfβ* and *Col3a1* expression in female SKO mice and *Mmp13* expression in male SKO mice (Fig. 3E, F). Moreover, there was a non-signficant trend toward a liraglutide-induced decrease in the expression of other markers of fibrosis examined (Fig. 3E, F). Liraglutide treatment did not alter expression of these genes in WT mice (data not shown). Thus liraglutide treatment significantly improved hepatic steatosis and liver fibrosis in female SKO mice with more modest improvements in male SKO mice.

**Figure 3:**
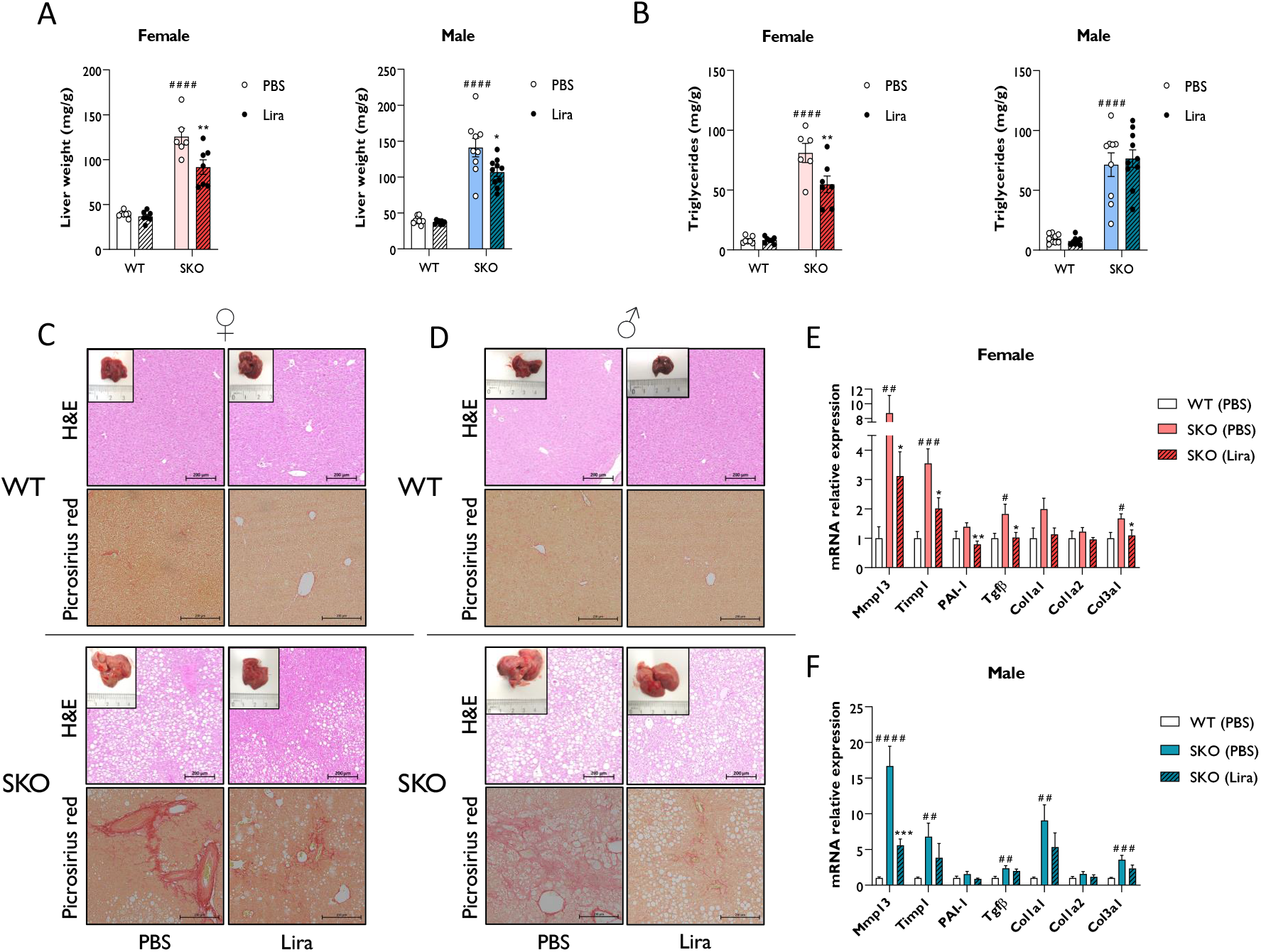
Liraglutide improves hepatic steatosis and fibrosis in mice with lipodystrophy. Male and female wild-type (WT) and seipin knockout (SKO) mice were intraperitoneally injected once daily with phosphate-buffered saline (PBS) or 0.2 mg/kg liraglutide (Lira) for 14 days. (**A**) Liver weight normalised to mice body weight. (**B**) Levels of hepatic triglycerides. (**C, D**) Representative images of hematoxylin and eosin (H&E) and picrosirius red-stained paraffin-embedded liver sections along with representative photographs of macroscopic liver appearance in female (**C**) and male (**D**) mice. Bars denote 200 µm. (**E, F**) Expression levels of a panel of fibrosis markers genes in WT mice and SKO mice treated with PBS or liraglutide. Gene expression was normalised to three reference genes (*Hprt, Nono* and *Ywhaz*) (female n= 6-7, male n= 8-10). Data are presented as mean ± SEM; *p < 0.05, **p < 0.01, ***p < 0.001 (PBS versus liraglutide within same genotype); ^#^p < 0.05, ^##^p < 0.01, ^###^p < 0.001, ^####^p < 0.0001 (WT (PBS) versus SKO (PBS)).

### 3.4 Liraglutide promotes insulin secretion in lipodystrophic mice

GLP-1 agonists have been shown to promote insulin secretion, β cell survival and β cell proliferation [10; 11; 24-26]. Fasting circulating insulin levels were very significantly higher in male and female SKO mice compared to WT mice and further increased by daily injections with liraglutide for 14 days (Fig. 4A). Insulin staining and detailed morphometric analysis of pancreatic sections revealed enlarged islets and increased pancreatic β cell area to total pancreatic area in SKO mice *versus* WT mice (Fig. 4B, C). Liraglutide- and PBS-treated SKO mice displayed comparable density of β cell area. However, in female but not male SKO mice liraglutide treatment significantly increased pancreas weight leading to an increase in total β cell mass per animal when normalised to body weight (Fig. 4D, E). To assess glucose stimulated insulin release (GSIS) blood glucose and insulin levels were determined before and atier administration of PBS or liraglutide followed by injection with D-glucose. SKO mice exhibited fasting blood glucose comparable to their WT counterparts, whilst fasting insulin levels were higher in SKO than in WT mice (Fig. 4F, G; Table S1). Liraglutide had no effect on glucose levels (Fig. 4F; Table S1) but increased plasma insulin concentrations in SKO mice, prior to glucose injection (Fig. 4G; Table S1). In response to a glucose bolus, blood glucose levels rapidly increased in PBS-treated WT and SKO mice with minimal fluctuation in plasma insulin levels (Fig. 4F, G; Table S1). Liraglutide markedly reduced these glycaemic excursions in both WT and SKO mice, and this occurred concomitantly with an increase in insulin secretion (Fig. 4G; Table S1). Although variable between mice, glucose stimulated insulin release was approximately 8-fold higher in liraglutide-treated than in PBS-treated SKO mice and 40 times higher than in liraglutide-treated WT mice. Thus, liraglutide significantly enhances the capacity of islets of SKO mice to secrete insulin in response to glucose.

**Figure 4:**
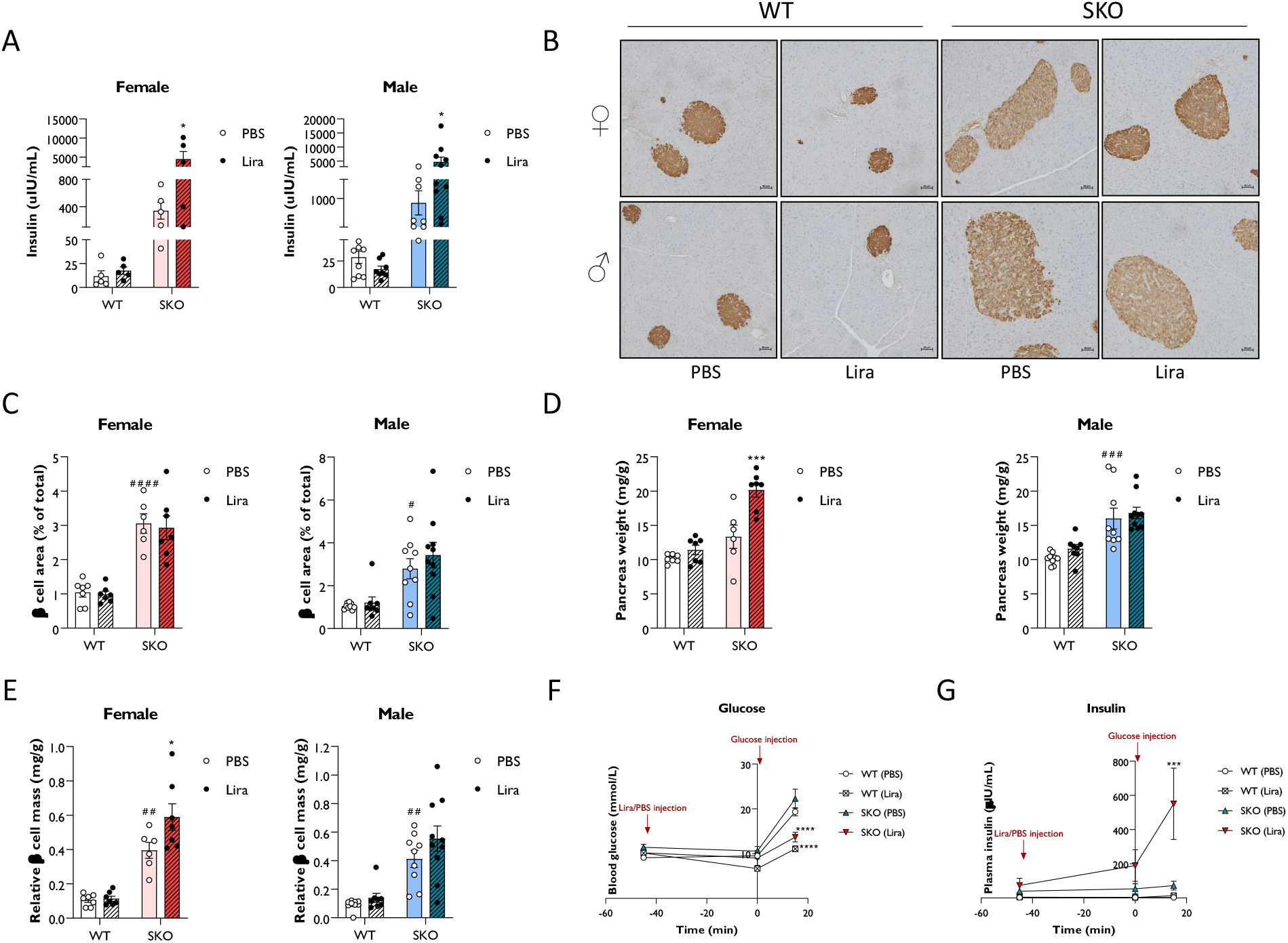
Liraglutide promotes insulin secretion in mice with lipodystrophy. Wild-type (WT) and seipin knockout (SKO) female and male mice were treated daily with 0.2 mg/kg liraglutide (Lira) for 14 days. Phosphate-buffered saline (PBS) was used as a vehicle control. (**A**) Effects of daily injections of liraglutide on serum insulin levels in female and male WT and SKO mice. (**B**) Representative images of insulin staining in pancreatic sections. (**C**) Quantification of insulin-stained pancreatic sections. Results are expressed as a percentage of the total pancreatic area occupied by β cells. (**D**) Pancreas weight normalised to body weight. (**E**) Relative β cell mass to pancreas weight normalised to body weight (female n= 5-7, male n= 8-10). (**F, G**) Acute effects of a single injection of 0.2 mg/kg liraglutide on both blood glucose (**F**) and plasma insulin (**G**) in male and female WT and SKO mice (n= 3-8). Data are presented as mean ± SEM; *p < 0.05, ***p < 0.001, ****p < 0.0001 (PBS versus liraglutide within same genotype); ^#^p < 0.05, ^##^p < 0.01, ^###^p < 0.001, ^####^p < 0.0001 (WT (PBS) versus SKO (PBS)).

## Discussion

Recombinant leptin therapy is currently the only dedicated therapy for generalised lipodystrophy but is effective only in some cases of partial lipodystrophy, making new therapies a clinical imperative [3; 5; 27-30]. Although GLP-1R agonists are widely used for common forms of type 2 diabetes, there are only three case study reports of their use and very few insights regarding their effects in patients with lipodystrophy [12-14].

Our data indicate that liraglutide enhances both insulin sensitivity and glucose regulated insulin release to alleviate lipoatrophic diabetes in SKO mice. In the liver, liraglutide treatment reduced hepatomegaly, steatosis, markers of fibrosis with evidence of acute improvements of gluconeogenesis from PTT assays. Faty liver disease in lipodystrophy is already acknowledged as distinct from other forms [31]. The work presented provides important new insights regarding exactly how faty liver disease in lipodystrophy may respond to GLP-1R agonist treatment. Given the severe lipodystrophy in our mouse model, our data also reveal that GLP-1R agonists can improve insulin sensitivity and metabolic health independent of adipose tissue, providing additional novel mechanistic insight.

Liraglutide induced only modest reductions in food intake in SKO mice, statistically significant only in males. This could be related to very low leptin levels in this model of CGL as previous studies have reported litle effect of GLP-1R agonists on food intake in mice and rats lacking functional leptin signalling [32; 33]. It may be that GLP-1R agonists may be most effective at reducing appetite in FPL patients with normal leptin levels, particularly valuable for this metreleptin-insensitive group. If so, the leptin requirement in CGL and FPL patients with low leptin levels could be reduced if used in combination with GLP-1R agonists. Similarly, the increased insulin secretion in our liraglutide treated SKO mice indicate that GLP-1R agonists could reduce the requirements for high dose insulin injections that are otien used by these individuals. Moreover, long acting and oral GLP-1R agonists could reduce or eliminate the considerable pain associated with insulin or leptin injections in these patients and therefore significantly improve quality of life for patients.

One potential concern for GLP-1R agonist use is the lipodystrophy-associated risk of pancreatis, which has also been suggested as a potential adverse effect of GLP-1R agonists. However, increased pancreatis has not been observed in long-term studies of liraglutide in rodents and primates [34-36]. Moreover, randomised controlled trials have not shown increased incidence of pancreatis in patients with type 2 diabetes treated with GLP-1R agonists [37-39]. Nonetheless, appropriate monitoring will be necessary for GLP-1R agonist use in the treatment of patients with lipodystrophy.

Our study suggests that GLP-1R agonists improve insulin sensitivity, glycaemic control, steatosis, and pancreatic beta cell function in CGL and, likely, in lipodystrophies more broadly. As the first study to examine in detail the effects of GLP-1R agonists in a pre-clinical model of lipodystrophy, we provide a valuable basis for future patient studies to ultimately relieve the suffering of individuals with lipodystrophy.

## ACKNOWLEDGEMENTS

The authors would like to thank the staff at the University of Aberdeen’s Microscopy and Histology Core Facility, the Medical Research Facility and Prof. Mirela Delibegovic for technical support and advice with the *in vivo* studies. This research was supported by funding from Diabetes UK (18/0005884 to JJR, RD Lawrence Fellowship 21/0006280 to GDM), the Biotechnology and Biological Sciences Research Council (BB/V015869/1 to JJR, BB/R01857X/1 and BB/V016849/1 to LKH), the Wellcome Trust Institutional Strategic Support Fund to the University of Aberdeen (to LKH and JJR). NS is supported by a BBSRC East of Scotland Bioscience Doctoral Training Partnership (eastBIO) PhD studentship. Research in the laboratory of WH is supported by Agency for Science, Technology and Research (A*STAR) Intramural funding, the Strategic Research Program (the Brain-Body Initiative, #21718), the Central Research Fund, and the JCO-VIP award.

## AUTHOR CONTRIBUTIONS

AR and JJR conceived the study and designed the experiments with input from LKH and GDM. AR, GDM and NS performed experiments and/or data analysis. WH provided reagents and intellectual contributions. AR and JJR wrote the manuscript with input from all other authors.

