## Supplementary tables for "GLP-1 Receptor Agonist Improves Metabolic Disease in a Pre-clinical Model of Lipodystrophy"

**Supplementary materials**

|  | **Glucose (mmol/L)** | | | **Insulin (μIU/mL)** | | |
| --- | --- | --- | --- | --- | --- | --- |
|  | ***- 45 min*** | ***0 min*** | ***+ 15 min*** | ***- 45 min*** | ***0 min*** | ***+ 15 min*** |
| **WT (PBS)** | 9.08 ± 0.44 | 9.51 ± 0.69 | 19.34 ± 0.90 | 3.19 ± 1.16 | 3.06 ± 0.65 | 2.93 ± 0.35 |
| **WT (Lira)** | 10.13 ± 0.43 | 6.65 ± 0.52 | 11.05 ± 0.66 | 5.33 ± 1.48 | 3.16 ± 1.32 | 13.59 ± 4.50 |
| **SKO (PBS)** | 11.42 ± 0.69 | 10.56 ± 1.03 | 22.25 ± 2.13 | 39.77 ± 9.68 | 53.46 ± 29.26 | 72.05 ± 25.41 |
| **SKO (Lira)** | 10.18 ± 0.95 | 9 ± 1.63 | 13.68 ± 1.09 | 73.91 ± 41.79 | 189.88 ± 92.66 | 550.74^***^ ± 208.75 |

**Table S1. Glucose-stimulated assay *in vivo*.** Acute effects of a single injection of 0.2 mg/kg liraglutide on both blood glucose and plasma insulin levels in male and female WT and SKO mice (female n= 6-7, male n= 8-10). Data are presented as mean ± SEM; *p < 0.05, ***p < 0.001, ****p < 0.0001 (*PBS versus Lira within same genotype; ^#^WT (PBS) versus SKO (PBS)).

| **Gene** | **Forward primer (5’ - 3’)** | **Reverse primer (5’ - 3’)** |
| --- | --- | --- |
| ***Col1a1*** | GCTCCTCTTAGGGGCCACT | CCACGTCTCACCATTGGGG |
| ***Col1a2*** | AAGGGTGCTACTGGACTCCC | TTGTTACCGGATTCTCCTTTGG |
| ***Col3a1*** | CCTGGCTCAAATGGCTCAC | CAGGACTGCCGTTATTCCCG |
| ***Hprt*** | GTTAAGCAGTACAGCCCCAAA | AGGGCATATCCAACAACAAACTT |
| ***Mmp13*** | TGTTTGCAGAGCACTACTTGAA | CAGTCACCTCTAAGCCAAAGAAA |
| ***Nono*** | GCCAGAATGAAGGCTTGACTAT | TATCAGGGGGAAGATTGCCCA |
| ***PAI-1*** | CAAGCTCTTCCAGACTATGGTG | ACCTTTGGTATGCCTTTCCAC |
| ***Tgfβ*** | CTCCCGTGGCTTCTAGTGC | GCCTTAGTTTGGACAGGATCTG |
| ***Timp1*** | CGAGACCACCTTATACCAGCG | ATGACTGGGGTGTAGGCGTA |
| ***Yhwaz*** | GAAAAGTTCTTGATCCCCAATGC | TGTGACTGGTCCACAATTCCTT |

**Table S2. Primers and sequences for real-time PCR assays.**
